# The phloem-resident OCTOPUS protein is a novel regulator of flg22-induced responses in *Arabidopsis thaliana*

**DOI:** 10.1101/2022.01.27.478095

**Authors:** Kaitlyn N. Greenwood, Courtney L. King, Isabella Melena, Katherine A. Stegemann, Carina A. Collins

## Abstract

Phloem is a critical tissue that transports photosynthates and extracellular signals in vascular plants. Although a functional phloem is necessary for plant health, it is also an ideal environment for pathogens to access host nutrients to promote pathogenesis. Even though many vascular pathogens induce economically relevant crop damage, very little is known about the mechanism(s) by which phloem cells detect potential pathogens and signal to minimize damage. Our lab searched existing phosphoproteomic databases, mining for proteins that were phosphorylated in response to the defense-elicitor flagellin, or flg22, AND were expressed in vascular cells, and we identified Octopus (OPS). OPS is polarly associated with the plasma membrane (PM) of sieve element cells and promotes their differentiation from procambial precursor cells by inhibiting the function of BIN2 in brassinosteroid-related signaling. The observation that OPS is differentially phosphorylated in response to flg22 led us to the examine whether OPS may function in flg22-induced signaling using *Arabidopsis* T-DNA insertion mutants lacking a functional *OPS*. In wild-type (WT) seedlings, flg22 binds to the PM receptor flagellin sensing 2 (FLS2) to initiate three branches of a signaling cascade that culminates in increased expression of distinct marker genes. Ultimately these signaling pathways lead to the restriction of pathogen growth. Two independent alleles of *ops* were treated with 100 *μ*M flg22 and marker genes from all three branches of FLS2 signaling exhibited higher expression than WT. We also found that in the absence of any flg22, *ops* mutants displayed increased flg22 signaling responses. Our results indicate that OPS may function as a negative regulator of flg22-induced signaling events and is one of very few phloem-resident proteins with a documented role in flg22 signaling. These results indicate that the phloem may be able to sense and response to the threat of bacterial pathogens in a unique way.

## Introduction

The phloem is a critical tissue in vascular plants that transports photosynthetic sugars from source tissues (leaves) to sink tissues (roots, flowers, and fruit); but this carbon-rich nutrient content also makes the phloem an ideal tissue for pathogenic microbes to colonize (1). There are many examples of phloem-limited bacterial pathogens, including the devastating citrus pathogen *Candidatus* Liberibacter asiaticus (CLas), the causal agent of citrus-greening disease (or ‘Huanglongbing’). CLas gains access to the phloem of citrus trees after being deposited by the brown planthopper or related insect vectors (2, 3). The phloem is such a rich source of nutrients that some pathogens that do not colonize the phloem directly will stimulate the aberrant development of phloem cells in other tissues to tap its resources (4). Two distinct cell types comprise the phloem transport system in plants: sieve elements and companion cells. Sieve elements are elongated cells whose primary function is to transport sap contents (photosynthetic sugars, RNA, peptides, and other small organic molecules) from source to sink. Adjacent to sieve elements are the companion cells, which are responsible for loading sap contents into sieve elements (5).

Current models of immune signaling in response to a pathogen generally assume that all host cells must individually recognize extracellular pathogen-associated molecular patterns (PAMPs). PAMPs are detected by plasma membrane (PM)- localized pattern recognition receptor (PRR) proteins that, after PAMP binding, can initiate intracellular signaling and activate robust defenses (6, 7). The classic example of a PRR is FLAGELLIN SENSING 2 (FLS2), a member of the leucine-rich repeat receptor-like kinase (LRR-RLK) family of receptors. FLS2 binds to the bacterial motor protein flagellin or to a conserved 22-amino acid peptide, flg22, derived from bacterial flagellin, to initiate a cascade of intracellular signaling events that contribute to the host immune response (8). Interestingly, FLS2 signaling events fall into one of three branches, each of which culminates in expression of marker genes (9, 10). For example, the calcium-dependent branch of FLS2 signaling induces expression of *PHOSPHATE-INDUCIBLE 1 (PHI1*), the mitogen-activated protein kinase (MAPK) pathway activates *FLG22-INDUCED RECEPTOR KINASE* (*FRK1*), and activation of the salicylic acid (SA) pathway causes increased expression of *PATHOGENESIS-RELATED1* (*PR1*) (7, 11, 12). In addition to changes in gene expression, FLS2 also initiates the production of extracellular reactive oxygen species (ROS) and the deposition of callose at the cell wall (7, 11, 12). Combined, these independent signaling events promote the restriction of pathogen growth and promote host immunity (12).

Largescale phosphoproteomic screens have been used to detect proteins that are differentially phosphorylated in response to flg22, potentially identifying proteins involved in the regulation of flg22-induced signaling (9, 13, 14). To find potential regulators of flg22-induced signaling in the phloem, we mined existing flg22 phosphoproteomic datasets and identified OCTOPUS (OPS), a protein that is differentially phosphorylated in response to flg22 in *Arabidopsis* suspension cell cultures (15). This finding was interesting because to the extent that its function is known, previous data indicates that OPS has a role in phloem development, not immune signaling (16–20). *Arabidopsis ops* mutant plants, which lack OPS, display decreased phloem pattern complexity, and contain undifferentiated sieve element cells, resulting in gaps in phloem strands of the root (16–18, 21). Further investigations indicated that OPS may regulate phloem developmental processes by promoting signaling events after perception of the hormone brassinolide (BL), a member of the brassinosteriod (BR) class (22). BL is an endogenous steroid hormone detected by the BRassinosteroid Insensitive-1 (BRI1) receptor kinase. When BRI1 binds BL, a series of downstream signaling events occurs, leading to cell elongation and growth. OPS interacts with a member of the GLYGOGEN SYNTHASE KINASE3 (GSK3) family, Brassinosteroid Insensitive-2 (BIN2), at the PM (22). The retention of BIN2 at the PM prevents BIN2 from inhibiting transcriptional changes needed to induce cell growth. Using #-glucuronidase (GUS) and GFP to visualize expression of OPS in *Arabidopsis*, these previous OPS studies found that OPS is present only in the phloem (16). Combined, this evidence indicates that OPS is an ideal candidate for studying flg22-signaling in the phloem.

Despite the devastating effects of phloem-dwelling pathogens on crops, it is not yet known in detail how the cells of the phloem detect the presence of bacterial pathogens or how they respond to them (1). Identifying the contributions of the phloem to pathogen detection and response will increase our knowledge of the tissue-specific mechanisms immune signaling. Using loss-of-function T-DNA mutants and gene expression analysis, we identify the sieve element protein OCTOPUS (OPS) as a regulator of flg22-induced signaling events in *Arabidopsis* seedlings. To our knowledge, this is the first example of a protein localized in sieve elements with such a role in flg22 signaling.

## Results

### Root length is reduced in ops-3 and ops-4 seedlings

To explore a potential role for OPS in flg22-elicited immune signaling, we obtained previously described T-DNA insertion mutant lines *ops-3* and *ops-4* (20). Using qPCR, we confirmed that both lines exhibit dramatically reduced *OPS* transcript levels (Fig. 1A-B). Consistent with previous studies (16), seedlings from both *ops* T-DNA lines displayed shorter roots with an increased amount of branching (Fig. 1 C-D). Despite the root growth defects, no gross morphological phenotypes, particularly in aerial tissue, were observed in either mutant as compared to Col-0.

**Figure 1.**
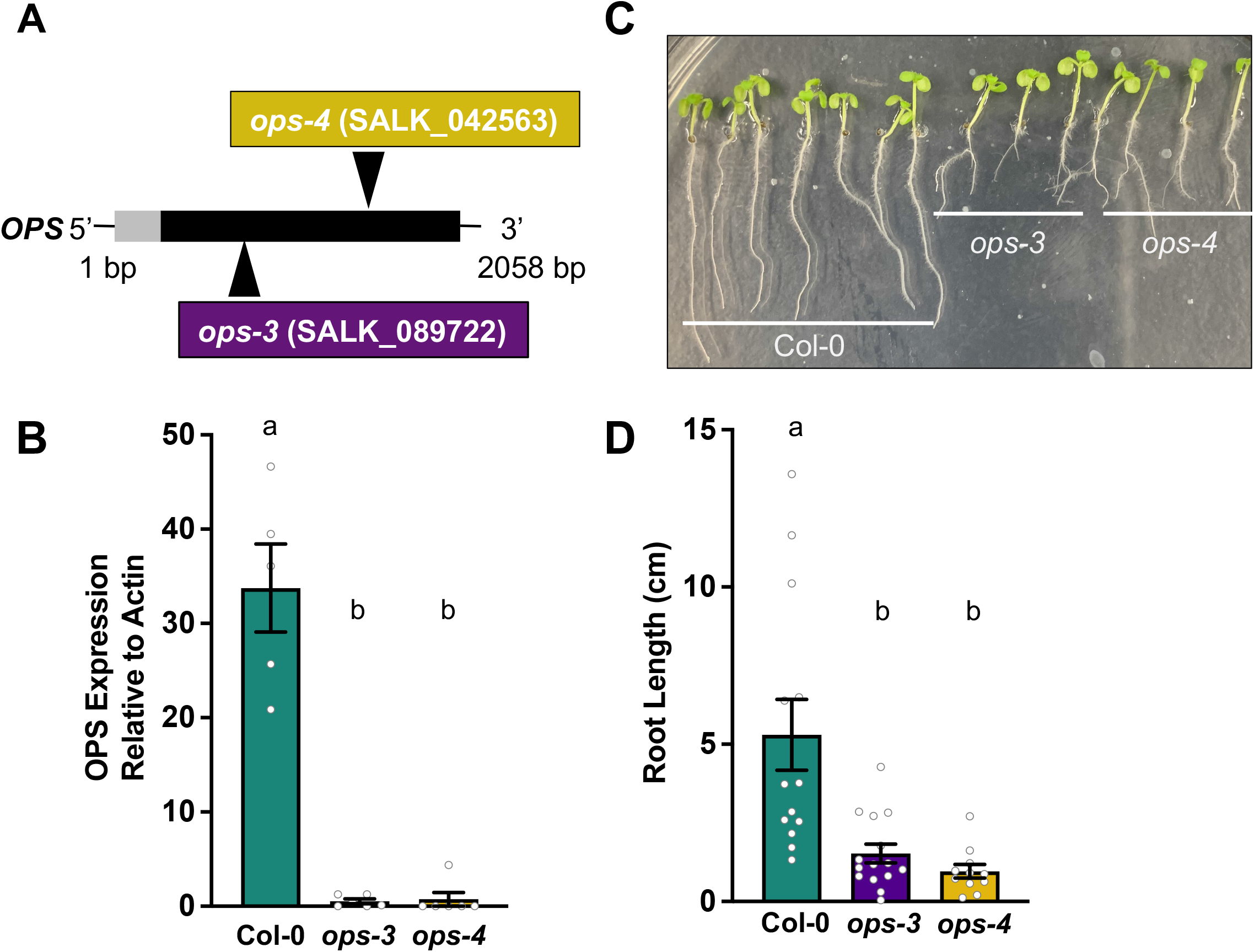
Characterization of two independent *OPS* T-DNA insertion lines. **A**. Location of T-DNA insertions in *OPS* and mutants used in this study. 20 **B**. Expression of *OPS* mRNA in Col0 and *ops* mutants measured using qPCR. *OPS* expression is shown relative to *Actin*, n=6. **C**. Comparison of root length between Col-0 WT, *ops-3* and *ops-4* mutants from vertically grown, day 7 old seedlings. Shown is a representative image. **D**. Analysis of root length in Col-0 WT, *ops-3*, and *ops-4* mutants. Measurements were made from images (as in C) using ImageJ, n = 20. Letters above all bars represent statistical differences as measured by a one-way ANOVA with Tukey multiple comparison test where *P < 0.05*. All error bars represent ±SE.

### flg22-induced responses are enhanced in ops-3 and ops-4 mutants

We hypothesized that because OPS was identified as a protein that was differentially phosphorylated after flg22 treatment in a phosphoproteomic screen (15), OPS might have a role in the regulation of flg22-induced signaling events in *Arabidopsis*. We first sought to determine how plants lacking OPS would respond to flg22 using a seedling growth inhibition assay. When Col-0 plants are grown in the presence of flg22 for an extended period (2 weeks), the seedlings experience growth stunting as a result of flg22 detection (8, 23). Col-0 and both independent *ops-3* and *ops-4* mutant alleles were grown in liquid media supplemented with either 0 or 1 *μ*M flg22 for two weeks before measuring their fresh weight. Growth of the Col-0 seedlings was inhibited by 74% (Fig. 2). Interestingly, both *ops* mutants showed significantly increased percent growth inhibition; 80% and 82%, respectively, when compared to Col-0 (Fig. 2). Importantly, the seedling growth inhibition was similar between *ops-3* and *ops-4*, indicating that this phenotype is indeed the result of loss of *OPS*.

**Figure 2.**
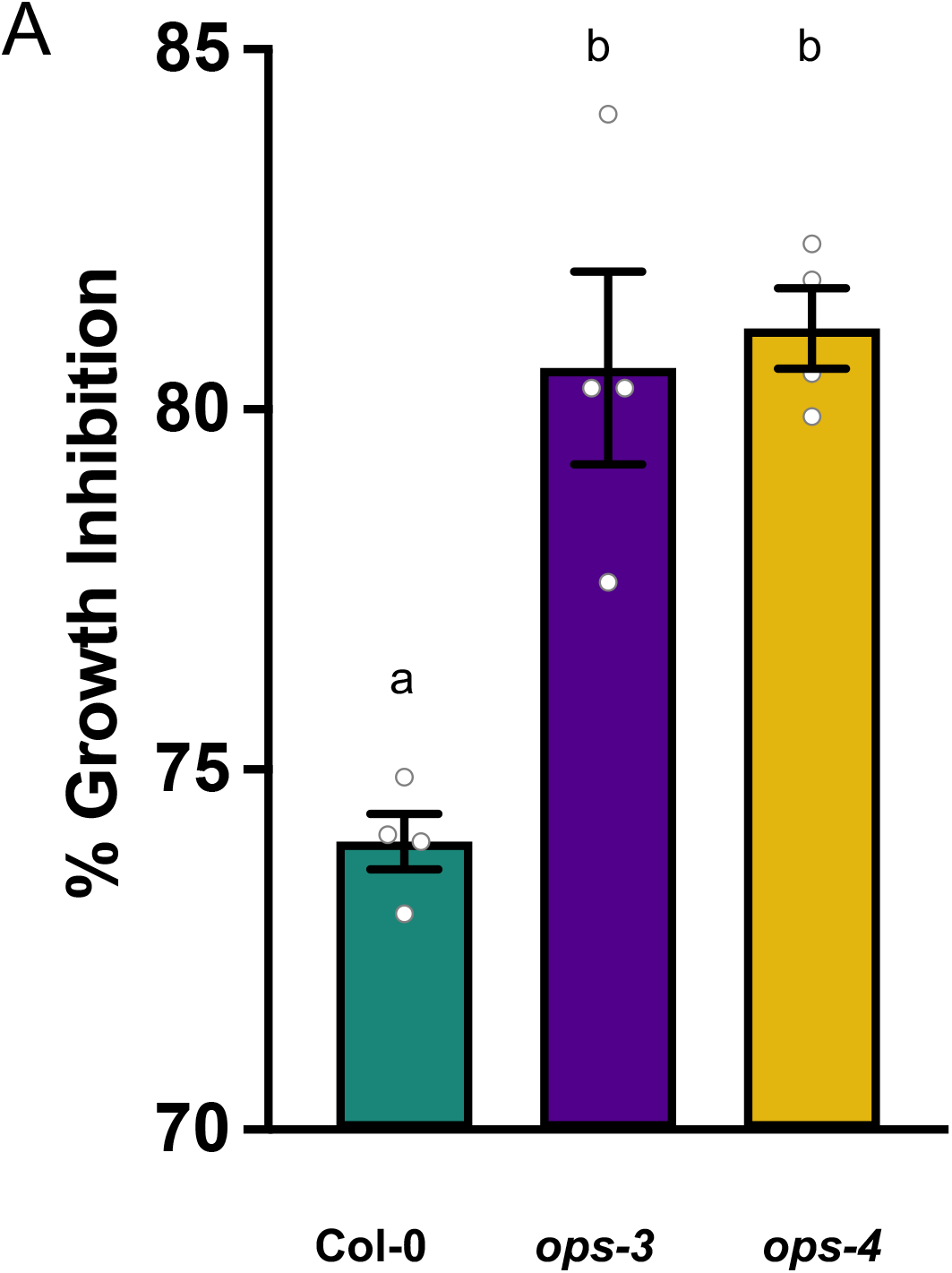
Lack of *OPS* increases flg22-induced growth inhibition. **A**. Four-day old seedlings were grown in the presence of 0 or 1 μM flg22 for two weeks before measuring their fresh weights and calculating the percent growth inhibition between growth conditions. Shown are percent change calculations pooled from 4 independent experiments, n = 40-60 seedlings per genotype per experiment. ANOVA with Tukey multiple comparison test with a *P* <0.05. All error bars are ±SE

Because we observed an increase of flg22-induced growth inhibition in *ops* mutants, which may be indicative of a defect in flg22 perceptions, we sought to investigate whether the lack of OPS led to a more specific flg22 signaling defect. In *Arabidopsis* when FLS2 binds flg22, a network of multiple signaling pathway branches are initiated, and each independent branch results in the expression of a pathway-specific marker gene (9, 10). To test if OPS has a role in one or more of these specific pathways, we measured induction of the flg22-induced and Ca^2+^ pathway-dependent expression of the marker gene *PHI1*. In Col-0 seedlings treated with 100 nM flg22, *PHI1* expression remains low prior to flg22 exposure (0 minutes), peaks within 30 minutes, and then falls to near base levels for 1-3 hours (Fig. 3A). When *ops-3* and *ops-4* seedlings were treated with 100 nM flg22 for the same time course, the expression pattern of *PHI1* was observably different. Even without the addition of 100 nM flg22, both *ops-3* and *ops-4* mutants showed significantly higher *PHI1* expression than Col-0 and while *PHI1* peaked in both mutants 30 minutes after treatment, expression levels were significantly higher than Col-0 (Fig. 3A). After observing that even in the absence of flg22, *ops* mutants had increased *PHI1* expression, we decided to examine subsequent marker genes in the absence of flg22.

**Figure 3.**
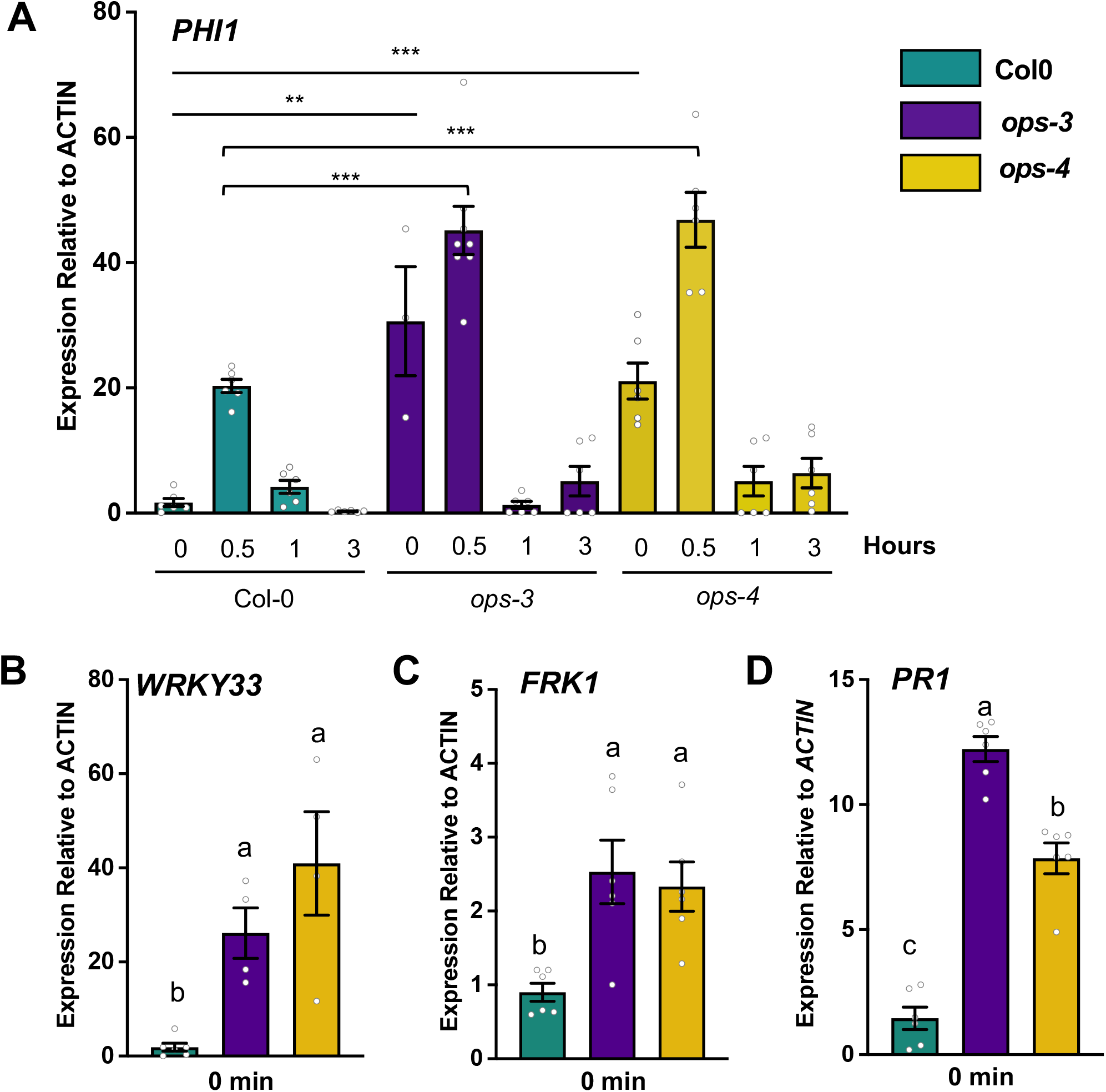
OPS functions as a negative regulator of flg22-induced immune signaling. **A**. Ten-day old old seedlings (Col-0 WT, *ops-3*, and *ops-4*) were treated with 100 nM flg22 and qPCR was used to measure expression of marker gene *PHI1* over a 3-hour time-course. Shown are means of pooled seedling samples (n = 4). Asterisks indicate significant differences compared with Col-0 0 min (lines) and Col-0 30 min (brackets) using a two-tailed Student’s *t*-test ** *P* > 0.01, *** *P* > 0.0001. **B-D**. qPCR analysis of untreated, ten-day old seedlings (Col-0 WT, *ops-3*, and *ops-4*). Expression of the marker genes *WRKY33* (**B**), *FRK1* (**C**), and *PR1* (**D**) was examined. Letters represent statistical differences as measured by one-way ANOVA with Tukey multiple comparison at *P <0.01*. All error bars are ±SE. Experiments were repeated at least three times with similar results.

MAPK activation after flg22 elicitation occurs independently of Ca^2+^ induction and leads to expression of *WRKY33 and FRK1*, so we next measured expression of the marker gene *FRK1*. Similarly to *PHI1*, *FRK1* expression was significantly higher in both *ops-3* and *ops-4* seedlings, even in the absence of flg22 (Fig. 3B-C). The final pathway we examined was the SA-dependent pathway of FLS2 signaling by measuring expression of the marker gene *PR1*. Consistent with our previous qPCR results, *PR1* expression was significantly increased in both *ops-3* and *ops-4* mutants. Together, these data suggest that OPS may be a negative regulator of flg22-induced immune responses.

### Brassinosteroid-induced hypocotyl growth is impaired in ops-3 and ops-4

Previous investigations of OPS function showed that OPS interacts with the kinase BRASSINOSTEROID-INSENSITIVE 2 (BIN2) at the PM, preventing BIN2 from inhibiting transcriptional changes needed to induce cell growth (22). These data indicate that OPS plays a role in promoting BL-induced cellular responses which may explain some of the phloem developmental defects observed in *ops* mutants (16, 24, 25); however, these observations were made by studying *Arabidopsis* plants expressing *OPS* under the control of constitutive promoter which may not accurately reflect *in planta* conditions.

To gain further evidence of a role for OPS in BL signaling, we treated Col-0 and *ops* mutants with 1 *μ*M epibrassinolide and measured hypocotyl elongation as a part of a standard BL- response assay. Col-0 seedlings exhibit elongated hypocotyls when exposed to BL, whereas elongation in both *ops-3* and *op-4* mutants was comparatively reduced. (Fig. 4A). When *Arabidopsis* seedlings are grown in the dark, their hypocotyls elongate to produce a BL-dependent etiolated phenotype (26). To gain further insights into the role of OPS in BR-mediated signaling, we measured the hypocotyls of Col-0, *ops-3* and *ops-4* seedlings grown in the light and the dark.

**Figure 4.**
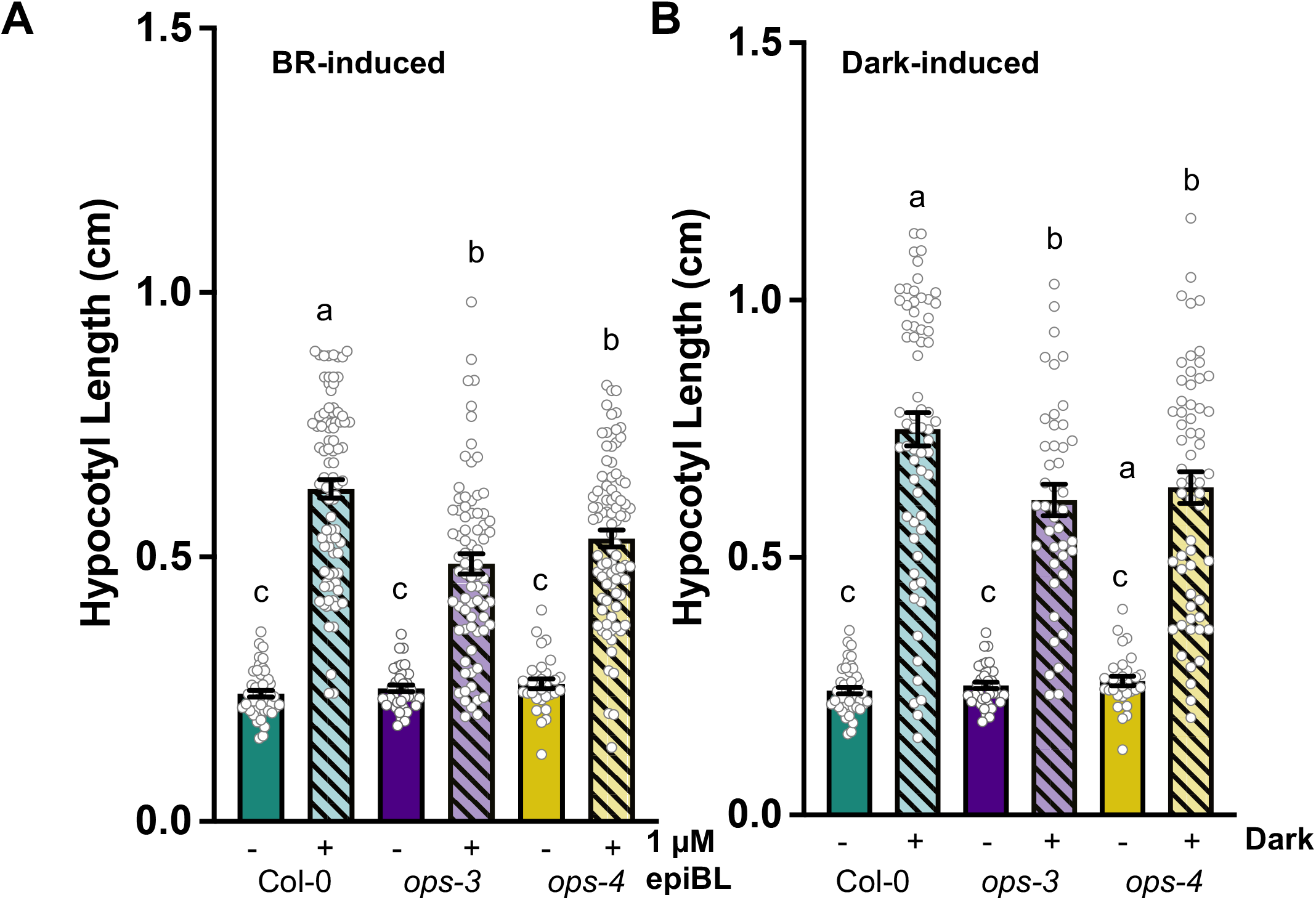
OPS is necessary for hypocotyl elongation. **A**. Hypocotyl length measurements taken of ten-day old seedlings (Col-0, ops-3, and *ops-4*) treated with 0 (−) or 1 (+) μM epibrassinolide. Values shown are pooled from three independent expriments, n = 60-90. **B**. Length of hypocotyls of ten-day old seedlings grown in the light (−) or the dark (+) for Col-0 WT, *ops-3*, and *ops-4*. Measurements are pooled from three independent experiments, n = 30-60. Letters represent statistical differences as measured by two-way ANOVA with Tukey multiple comparison test at *P* <0.05. Error bars are ±SE and all experiments were repeated at least three times with similar results.

Col-0 hypocotyls elongate significantly in the dark, and while the length of hypocotyls of both *ops-3* and *ops-4* seedlings do elongate, they are significantly shorter than those of Col-0 (Fig. 4B). These results provide additional evidence that OPS is required for BR-induced hypocotyl elongation.

## Discussion

Phloem-limited pathogens cause devastating crop losses, but our understanding of how vasculature tissue responds to pathogen invasion is incomplete, making it imperative to study the contributions of tissue-specific immune signaling to overall plant defenses. A new study demonstrated for the first time that when *Citrus sinesis* or “Valencia” trees are infected with the phloem-limited pathogen CLas they exhibit increased ROS production and callose deposition, indicative of a PAMP-triggered immune response (27). Moreover, authors showed that both sieve elements and companion cells underwent programmed cell death in response to prolonged CLas infection (27). Here, we have identified a protein exclusively expressed in sieve elements of the phloem that also has a role in flg22-induced immune responses, which we believe may be the first of its kind. These recent advances highlight the likelihood that phloem cells can detect the presence of bacterial pathogens and are active in signaling a response.

In this work, we demonstrate that two independent *Arabidopsis* lines harboring T-DNA insertions in the *OPS* gene lack *OPS* expression and display constitutive and increased flg22-induced expression of several immunity-related marker genes. Furthermore, we found that expression of these genes can be detected even in the absence of flg22. This combination of results indicates that OPS functions as a negative regulator of these flg22-induced responses and therefore OPS exhibits a suppressive effect on this pathway (Fig. 5). While the specific mechanism of OPS function in flg22 signaling remains unclear, OPS may suppress activation of *PHI1*, *FRK1*, *WRKY33*, and *PR1* expression by directly inhibiting a signaling event in the flg22 pathway. One possibility is OPS may be directly targeting another member of the GSK3 protein family that is instead known to regulate flg22-induced signaling, such as *Arabidopsis* Protein Kinase $ (ASK $) (28). Because genes from three independent branches of FLS2-flg22 signaling show the same increased expression in the *ops* mutant backgrounds, it is likely that if OPS is acting directly to regulate these signals, it must function early in flg22 signaling. A second possibility remains that OPS has an indirect role in flg22 signaling. OPS is expressed in the sieve elements of the phloem in *Arabidopsis* and previous studies identified that *ops* mutants display in incomplete sieve element differentiation which results in a discontinuous protophloem and metaphloem cell file in the roots (16, 17, 24). Therefore, our observation that *ops* mutants exhibit increased flg22 marker gene expression could be the result of incomplete phloem transport, resulting in the loss of transport of a yet unknown phloem-mobile regulator of flg22 responses.

**Figure 5.**
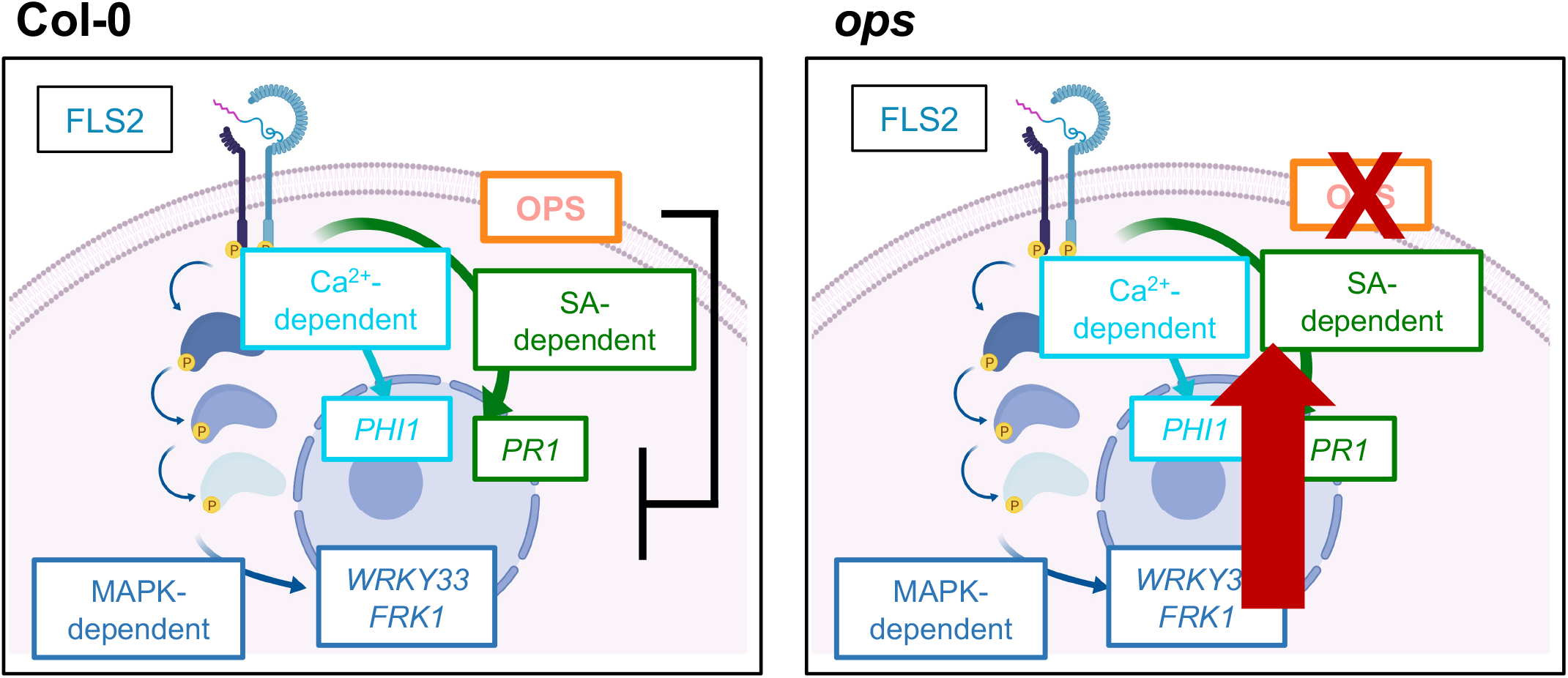
Summary of OPS flg22 signaling phenotype. Loss of the protein OPS (*ops;* right panel) results in increased expression of marker genes from three independent branches of the FLS2 signaling network (red arrow). This indicates that OPS plays a role in suppressing flg22-induced signaling (left panel).

Our results are consistent with reports that FLS2 expression is detected in the vasculature of *Arabidopsis* cotyledons in roots (29, 30), however detection of FLS2 could only be confirmed in the stele, which comprises both sieve elements and companion cells. FLS2 expression in the stele suggests that some cells of the vasculature are capable of detecting and responding to flg22, and while it has yet to be determined whether FLS2 expressed in the vasculature induces flg22 signaling events, our results indicate that some element of flg22 signaling does occur in the phloem.

In addition to our data demonstrating a role for OPS in flg22 signaling, we found that *Arabidopsis* loss-of-function *ops* mutants have shorter hypocotyls in response to BL treatments and in the dark. This is consistent with other work showing that OPS overexpression lines display increased BR responses (22), and confirms that OPS promotes BR-induced signaling for hypocotyl elongation. Finding that OPS functions in both the BRI1 and FLS2 signaling pathways is particularly interesting because these signaling pathways intersect downstream of initial receptor-ligand perception. Initiation of the BRI1 signaling pathway induces expression of several *WRKY* family transcription factors that inhibit the activation of gene expression of some flg22-inudced genes (none that were tested in this work) resulting in a suppression of defense responses (31, 32). That OPS functions as a positive regulator of BL signaling may explain why in *ops* mutants, we observe increased flg22-induced immune responses. While there is much left to understand about the function of OPS, it remains one of the few phloem-localized proteins identified with a role in immunity-related signaling events. Because it remains unknown whether sieve elements or companion cells can directly test and respond to PAMPs, future studies could use OPS as a model to begin answering these questions.

## Methods

### Plant Material and Growth Conditions

T-DNA insertion lines of Arabidopsis (*Arabidopsis thaliana*) *ops-3* SALK_089722 (20) and *ops-4* SALK_042563 (20) were obtained from the Arabidopsis Biological Resource Center at The Ohio State University (https://abrc.osu.edu/). Genotypes were confirmed using PCR (primers listed in Supplemental Table 1). Surface-sterilized seeds were sown on 0.5X Murashige and Skoog medium + 1% (w/v) sucrose solidified with 0.6% (w/v) agar as described. After 2 days stratification at 4°C, seedlings were germinated and grown at 22°C with an 10-h-light/14-h-dark photoperiod at 82 mmol m22 s21. Unless otherwise noted, seedlings were grown under these conditions for 10 days before sample analysis.

### Peptides and Hormones

The peptide flg22 (QRLSTGSRINSAKDDAAGLQIA) was purchased from Genscript and used for elicitation at the indicated concentrations and for the indicated times as described. 24-epibrassinolide (BL) was purchased from MilliporeSigma and used for the indicated times and concentrations described.

### Root and Hypocotyl Measurements

Root length measurements were performed as described (33). 10-day-old seedlings grown under the conditions described, were traced using Fiji Free-hand tool. For hypocotyl measurements, 4 day-old-seedlings were treated with the indicated concentration of BL in liquid media and placed at 22°C for an additional 7 days. Seedlings were removed and hypocotyls were traced using the Fiji Free-hand tool.

### Flg22 Seedling Growth Inhibition

Seedling growth inhibition was measured as previously described (23). Briefly, four-day-old seedlings were aseptically transferred from MS agar to wells of a 12-well microtiter plate (three seedlings per well) containing 1 ml of liquid MS medium with or without 1 *μ*M flg22. After 14 days, seedling fresh weights were recorded.

### RNA Isolation and RT-qPCR

RNA isolation, cDNA synthesis, and RT-qPCR were performed as previously described (34). Unless stated otherwise, for each sample, three to five seedlings were elicited with the indicated concentration of flg22 peptide and placed at 22°C for the time indicated. Tissue was flash frozen in liquid nitrogen at the indicated time points. Total RNA was isolated from tissue using Trizol reagent (Sigma-Aldrich) according to the manufacturer’s protocol. RT-qPCR was performed on cDNA using a Rotor-Q Real-Time PCR Cycler from Qiagen using gene-specific primers and normalized to the *ACTIN* gene (listed in Supplemental Table 1).

## References

1. Y. Jiang, C. X. Zhang, R. Chen, S. Y. He, Challenging battles of plants with phloem-feeding insects and prokaryotic pathogens. Proc Natl Acad Sci U S A 116, 23390–23397 (2019).

2. E. D. Ammar, J. George, K. Sturgeon, L. L. Stelinski, R. G. Shatters, Asian citrus psyllid adults inoculate huanglongbing bacterium more efficiently than nymphs when this bacterium is acquired by early instar nymphs. Sci Rep 10, 18244 (2020).

3. M. Ghanim et al., ’Candidatus Liberibacter asiaticus’ Accumulates inside Endoplasmic Reticulum Associated Vacuoles in the Gut Cells of Diaphorina citri. Sci Rep 7, 16945 (2017).

4. P. Walerowski et al., Clubroot Disease Stimulates Early Steps of Phloem Differentiation and Recruits SWEET Sucrose Transporters within Developing Galls. Plant Cell 30, 3058–3073 (2018).

5. R. Breia et al., Plant SWEETs: from sugar transport to plant-pathogen interaction and more unexpected physiological roles. Plant Physiol 186, 836–852 (2021).

6. D. Couto, C. Zipfel, Regulation of pattern recognition receptor signalling in plants. Nat Rev Immunol 16, 537–552 (2016).

7. V. Nicaise, M. Roux, C. Zipfel, Recent advances in PAMP-triggered immunity against bacteria: pattern recognition receptors watch over and raise the alarm. Plant Physiol 150, 1638–1647 (2009).

8. L. Gomez-Gomez, T. Boller, FLS2: an LRR receptor-like kinase involved in the perception of the bacterial elicitor flagellin in Arabidopsis. Mol Cell 5, 1003–1011 (2000).

9. J. M. Smith et al., Loss of Arabidopsis thaliana Dynamin-Related Protein 2B reveals separation of innate immune signaling pathways. PLoS Pathog 10, e1004578 (2014).

10. D. A. Korasick et al., Novel functions of Stomatal Cytokinesis-Defective 1 (SCD1) in innate immune responses against bacteria. J Biol Chem 285, 23342–23350 (2010).

11. J. Monaghan, C. Zipfel, Plant pattern recognition receptor complexes at the plasma membrane. Curr Opin Plant Biol 15, 349–357 (2012).

12. T. A. DeFalco, C. Zipfel, Molecular mechanisms of early plant pattern-triggered immune signaling. Mol Cell 81, 4346 (2021).

13. M. Kalde, T. S. Nuhse, K. Findlay, S. C. Peck, The syntaxin SYP132 contributes to plant resistance against bacteria and secretion of pathogenesis-related protein 1. Proc Natl Acad Sci U S A 104, 11850–11855 (2007).

14. T. S. Nuhse, A. R. Bottrill, A. M. Jones, S. C. Peck, Quantitative phosphoproteomic analysis of plasma membrane proteins reveals regulatory mechanisms of plant innate immune responses. Plant J 51, 931–940 (2007).

15. J. J. Benschop et al., Quantitative phosphoproteomics of early elicitor signaling in Arabidopsis. Mol Cell Proteomics 6, 1198–1214 (2007).

16. E. Truernit, H. Bauby, K. Belcram, J. Barthelemy, J. C. Palauqui, OCTOPUS, a polarly localised membrane-associated protein, regulates phloem differentiation entry in Arabidopsis thaliana. Development 139, 1306–1315 (2012).

17. M. A. Ruiz Sola et al., OCTOPUS-LIKE 2, a novel player in Arabidopsis root and vascular development, reveals a key role for OCTOPUS family genes in root metaphloem sieve tube differentiation. New Phytol 216, 1191–1204 (2017).

18. A. S. Breda, O. Hazak, C. S. Hardtke, Phosphosite charge rather than shootward localization determines OCTOPUS activity in root protophloem. Proc Natl Acad Sci U S A 114, E5721–E5730 (2017).

19. Y. H. Kang, A. Breda, C. S. Hardtke, Brassinosteroid signaling directs formative cell divisions and protophloem differentiation in Arabidopsis root meristems. Development 144, 272–280 (2017).

20. H. Roschzttardtz et al., The VASCULATURE COMPLEXITY AND CONNECTIVITY gene encodes a plant-specific protein required for embryo provasculature development. Plant Physiol 166, 889–902 (2014).

21. B. Gujas et al., A Reservoir of Pluripotent Phloem Cells Safeguards the Linear Developmental Trajectory of Protophloem Sieve Elements. Curr Biol 30, 755–766 e754 (2020).

22. P. Anne et al., OCTOPUS Negatively Regulates BIN2 to Control Phloem Differentiation in Arabidopsis thaliana. Curr Biol 25, 2584–2590 (2015).

23. J. C. Anderson et al., Arabidopsis MAP Kinase Phosphatase 1 (AtMKP1) negatively regulates MPK6-mediated PAMP responses and resistance against bacteria. Plant J 67, 258–268 (2011).

24. A. Rodriguez-Villalon et al., Molecular genetic framework for protophloem formation. Proc Natl Acad Sci U S A 111, 11551–11556 (2014).

25. A. Rodriguez-Villalon, B. Gujas, R. van Wijk, T. Munnik, C. S. Hardtke, Primary root protophloem differentiation requires balanced phosphatidylinositol-4,5-biphosphate levels and systemically affects root branching. Development 142, 1437–1446 (2015).

26. S. Fujioka et al., The Arabidopsis deetiolated2 mutant is blocked early in brassinosteroid biosynthesis. Plant Cell 9, 1951–1962 (1997).

27. W. Ma et al., Citrus Huanglongbing is a pathogen-triggered immune disease that can be mitigated with antioxidants and gibberellin. Nature Communications 13, 529 (2022).

28. H. Stampfl, M. Fritz, S. Dal Santo, C. Jonak, The GSK3/Shaggy-Like Kinase ASKalpha Contributes to Pattern-Triggered Immunity. Plant Physiol 171, 1366–1377 (2016).

29. M. Beck et al., Expression patterns of flagellin sensing 2 map to bacterial entry sites in plant shoots and roots. J Exp Bot 65, 6487–6498 (2014).

30. I. Wyrsch, A. Dominguez-Ferreras, N. Geldner, T. Boller, Tissue-specific FLAGELLIN-SENSING 2 (FLS2) expression in roots restores immune responses in Arabidopsis fls2 mutants. New Phytol 206, 774–784 (2015).

31. R. Lozano-Duran et al., The transcriptional regulator BZR1 mediates trade-off between plant innate immunity and growth. Elife 2, e00983 (2013).

32. Y. Belkhadir, L. Yang, J. Hetzel, J. L. Dangl, J. Chory, The growth-defense pivot: crisis management in plants mediated by LRR-RK surface receptors. Trends Biochem Sci 39, 447–456 (2014).

33. G. Ekanayake et al., DYNAMIN-RELATED PROTEIN DRP1A functions with DRP2B in plant growth, flg22-immune responses, and endocytosis. Plant Physiol 185, 1986–2002 (2021).

34. C. A. Collins et al., EPSIN1 Modulates the Plasma Membrane Abundance of FLAGELLIN SENSING2 for Effective Immune Responses. Plant Physiol 182, 1762–1775 (2020).

